# Distinct functional roles across hippocampal subfields in anticipation to event boundaries during schema learning

**DOI:** 10.64898/2025.12.01.691548

**Authors:** Silvy H.P. Collin

## Abstract

Continuous experiences are constantly segmented into separate events in memory, which is accompanied by increased hippocampal activity at the boundaries between these events. Extracting knowledge across all these experiences about what type of events to expect in a certain environment leads to event schemas being represented in the brain. These event schemas help to predict future events. What are the differences across the various distinct hippocampal subfields in peaks of activity around event boundaries? And what is the influence of event schema learning on these hippocampal subfield peaks? Here, this was investigated using data from an fMRI experiment in which participants learned two (related) event schemas by exposure to many similar animated videos of wedding ceremonies in a novel fictional culture. Results revealed that the hippocampus (CA1, CA2/3, dentate gyrus and subiculum) showed peaks in activity before event boundaries. Interestingly, the CA1, CA2/3 and dentate gyrus pre-boundary peaks attenuated due to event schema learning. This provides evidence for these subfields signaling uncertainty about upcoming events in the seconds prior to event boundaries. These results give a more precise understanding about the temporal dynamics around event boundaries in the distinct hippocampal subfields, and how these hippocampal subfields’ responses interact with schema learning.

## 1 Introduction

Event segmentation divides ongoing experience into meaningful units by generating event boundaries at points of change, and it was shown that these boundaries reliably evoke increases in hippocampal activity (Baldassano et al., 2017; Barnett et al., 2024; Ben-Yakov & Henson, 2018; Bilkey & Jensen, 2021; Brunec et al., 2018; Griffiths & Fuentemilla, 2020; Reagh et al., 2020; Zacks et al., 2001). Breaking our experiences into meaningful units in memory is essential for making sense of them (De Soares et al., 2024; Kurby & Zacks, 2008; Radvansky & Zacks, 2017; Zacks et al., 2007). Event segmentation is spontaneous and strongly influences how experiences are processed and remembered (Flores et al., 2017; Sargent et al., 2013; Sava-Segal et al., 2023). According to event segmentation theory, novel or unexpected occurrences generate prediction errors that trigger the creation of event boundaries (Zacks et al., 2007). Many studies have focused on the neural mechanisms behind this spontaneous segmentation process (Baldassano et al., 2017; Geerligs et al., 2021).

Ben-Yakov and Henson, 2018 showed that more salient boundaries evoke stronger hippocampal responses. At the same time, we develop event schemas that encode what events are typically expected in a given environment (Audrain & McAndrews, 2022; Baldassano et al., 2018; Bein & Niv, 2025; Beukers et al., 2023; Brod et al., 2015; Collin et al., 2025; DuBrow et al., 2017; Ghosh & Gilboa, 2014; Gilboa & Marlatte, 2017; Greve et al., 2019; Höltje et al., 2019; Masis-Obando et al., 2022; Roy et al., 2024; van Kesteren et al., 2010, 2012, 2020). By providing strong predictions about upcoming events, these event schemas may reduce the subjective salience of event transitions, thereby decreasing the likelihood of large prediction errors and potentially diminishing hippocampal boundary responses. Schema learning might also shift the timing of hippocampal boundary responses, as learning what to expect could lead to a forward shift in boundary-related activity (Michelmann et al., 2021).

Thus, while the function of the hippocampus in signaling event boundaries (Barnett et al., 2024; Brunec et al., 2018; Griffiths & Fuentemilla, 2020) and its relation to prediction (Ben-Yakov & Henson, 2018; Sinclair et al., 2021) is well established, knowledge about possible differences across the various hippocampal subfields (i.e., CA1, CA2, CA3, dentate gyrus, subiculum) for such increases in neural activity at event boundaries and its relation to schema learning is relatively sparse. Chen et al., 2024 uses a computational model to suggest that it is in particular the CA1 subfield that is sensitive to prediction error and relates this sensitivity to novelty detection. Few human neuroimaging studies have focused on this topic. Therefore, research questions for this project were: *How do the different hippocampal subfields respond in anticipation to event boundaries? And what is the influence of event schema learning on the peaks in neural activity in anticipation to event boundaries across these hippocampal subfields?*

Various studies have in general investigated the functional dissociation across these hippocampal subfields in memory. The subiculum is mostly involved in memory retrieval tasks while the other subfields are suggested to be involved in both memory retrieval and memory encoding tasks (Seok & Cheong, 2020). Many studies that focus on functional differentiation across hippocampal subfields focus on distinguishing them based on either pattern separation or pattern completion. Whether a certain situation triggers pattern separation or completion is believed to depend on how well the current environment aligns with past experiences. When the situation matches expectations, the hippocampus tends to support retrieval of an old representation through pattern completion. However, if there is enough of a mismatch, it switches to a mode that promotes encoding of the novel event, relying on pattern separation (Rolls, 2013). The CA3 has both been shown to be involved in integrating related information (Bein & Davachi, 2024; Bonnici et al., 2012; Dimsdale-Zucker et al., 2018; Farovik et al., 2010; Grande et al., 2019; S. Leutgeb & Leutgeb, 2007), and by other studies shown to differentiate between related information (Leal et al., 2014; Molitor et al., 2021). The CA1 is primarily implicated in memory integration, for example in pattern completion (Bonnici et al., 2012) and in integrating information over longer periods of time (Farovik et al., 2010). In contrast to this, dentate gyrus has rather consistently been shown to be involved in creating distinct representations of highly similar information in order to avoid interference in memory (Azab et al., 2014; Baker et al., 2016; Bakker et al., 2008; Bein & Davachi, 2024; Myers & Scharfman, 2011).

The prediction of the current study was that hippocampal subfields that were shown to be involved in memory encoding tasks (CA1, CA2/3, dentate gyrus) (Seok & Cheong, 2020) would show increases in hippocampal activity in anticipation to event boundaries. Furthermore, as a result of learning the new event schema, participants might start to integrate the at first seemingly distinct events due to learning of the underlying event schemas. CA1 and CA3 are both suggested to be involved in memory integration of related information (Bein & Davachi, 2024; Dimsdale-Zucker et al., 2018; Farovik et al., 2010; Grande et al., 2019; S. Leutgeb & Leutgeb, 2007; Molitor et al., 2021). For this reason, it can be expected that the magnitude of the expected CA1 peaks and CA3 peaks of activity at event boundaries would decrease as a result of event schema learning, since schema learning would lead to integrating of the events, which would lead to less novelty and uncertainty at the boundaries between the events within the sequences.

## 2 Methods

This study presents a re-analysis of functional MRI data originally reported by Collin et al., 2025 as collected at the Princeton Neuroscience Institute (Princeton University, USA). Forty participants (ages 18–35) took part in a two-day fMRI experiment featuring animated wedding videos from a fictional culture. Each video depicted a 2-minute ceremony with ritual sequences that differed depending on whether the couple belonged to the North or South of an imagined island. While the ritual patterns followed consistent trajectories (two possible sequences for North and two for South), participants were not explicitly told about these regional distinctions and instead had to infer them based on contextual cues (see Figure 1 in Collin et al., 2025 for the study design). Each ritual path included a series of four events: beginning with either a campfire or flower ritual, followed by a torch or coin ritual, then an egg or painting ritual, and ending with a gift ritual. On day 1, participants viewed wedding videos of the North for two blocks followed by wedding videos of the South for 2 blocks (or vice versa). On day 2, they viewed a new set of 12 wedding videos followed by a recall task. Participants provided informed consent and were compensated for their time (in accordance with experimental procedures approved by the Princeton University Institutional Review Board, IRB protocol 7883). Full details regarding stimuli, tasks, procedures, and MRI pre-processing are available in Collin et al., 2025. The current study focuses on analyzing day 1 of this project (while Collin et al., 2025 was mostly focused on analyzing the data of day 2 of this project). More specifically, in the current study the focus was on the hippocampal subfields during novel schema learning on day 1.

**Fig. 1.**
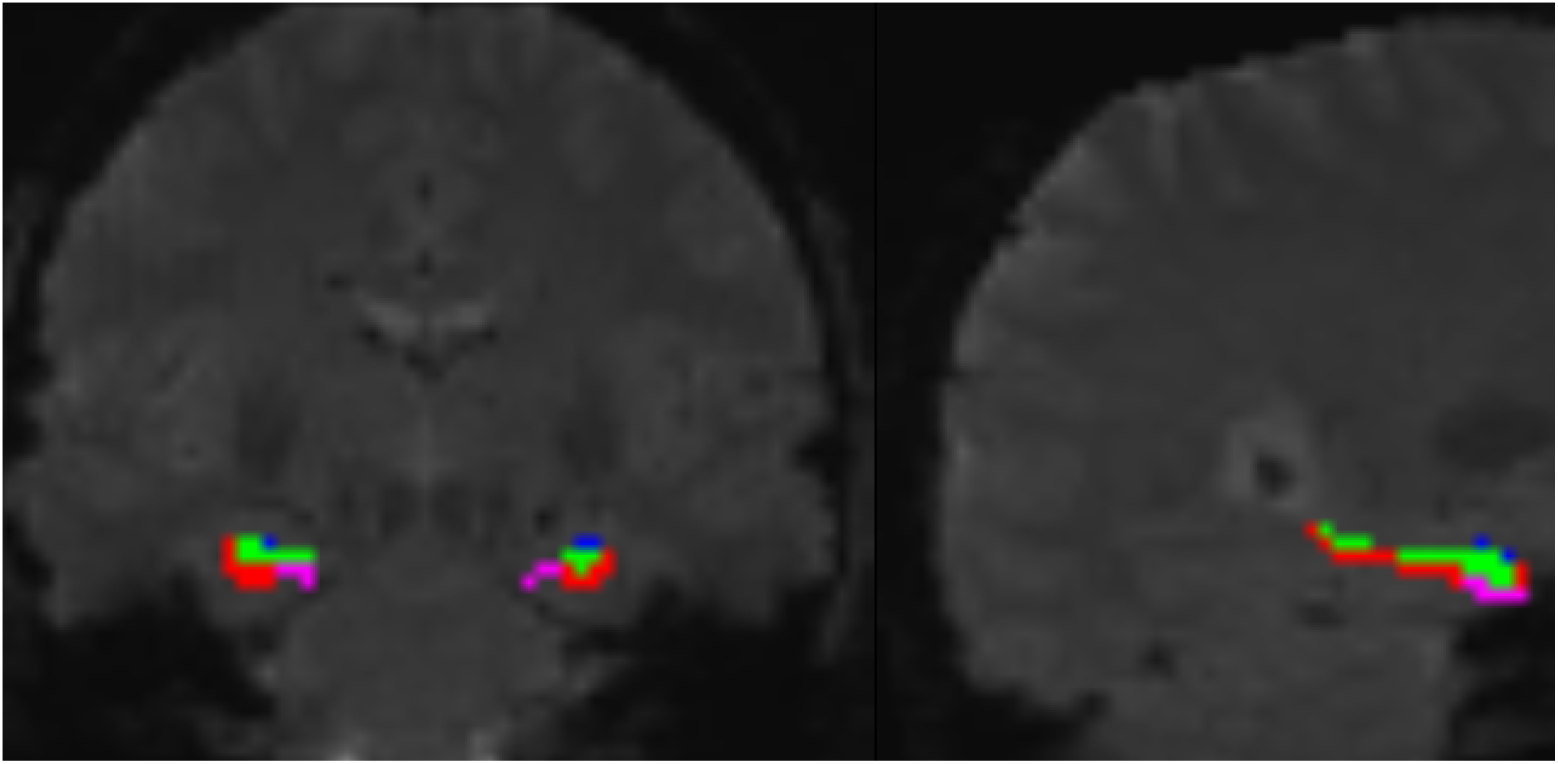
An example participants’ hippocampal subfield segmentation as performed by ASHS. Red is CA1, blue is CA2/3, green is dentate gyrus, pink is subiculum.

### 2.1 Hippocampal subfields

For the purpose of analyzing hippocampal subfields, an additional anatomical scan was performed (i.e., additional to the T1w-image). A T2-weighted turbo spin-echo (TSE) image was collected: TR = 11390 ms, TE = 90 ms, voxel size = 0.44 x 0.44 x 1.5 mm, 54 slices. The image was positioned to be acquired perpendicular to the long axis of the hippocampus. This TSE scan in combination with the T1w-images were used to determine the hippocampal subfields. For this purpose, the Automated Segmentation of Hippocampal Subfields (ASHS) toolbox (Yushkevich et al., 2015) was used. Each participant’s fMRIPrep-preprocessed T1w template and the TSE image were submitted as input to ASHS which led to subfield segmentations as output. Specifically, to a mask for the CA1, CA2/3, dentate gyrus (DG), and subiculum, separately for each hemisphere. These masks were then transformed to match the functional data. See Figure 1 for an example participants’ hippocampal subfield segmentation. All further described analyses of the hippocampal subfields were performed using the fMRI data in T1w (subject-specific) space.

### 2.2 Analyses

#### 2.2.1 Univariate activity around event boundaries

To test whether the hippocampus indeed showed increased neural activity around event boundaries, the fMRI data transformed to MNI152NLin2009cAsym space during each of the 4 schema learning blocks of day 1 was used. The MR images around event boundaries was cut out (from 5 MR images before an event boundary to 5 MR images after an event boundary, after shifting the entire timeline of images with 3 TRs (i.e., 4.5 sec) to account for the hemodynamic BOLD response), using the event boundaries between the various stages within any given wedding ceremony. Thus, event boundaries in this study were defined as the objective event boundaries (i.e., switches between the event videos within a wedding). To determine if and when the hippocampus showed a peak in activity in anticipation to event boundaries, a paired-samples t-test was used to compare the average activity in that anticipatory period (i.e., TR = -5 until TR = -1) which was tested against the average activity in the post-event boundary period (i.e., TR = 0 until TR = 4). This analysis was repeated for each hippocampal subfield.

Furthermore, the following analysis determined whether the peak in activity in anticipation to event boundaries was higher in magnitude in first schema learning blocks compared to second schema-learning blocks. A separate average of the TRs in the anticipatory period (i.e., TR = -5 until TR = -1) and of the TRs in the post-event boundary period (i.e., TR = 0 until TR = 4) for the two first blocks (i.e., block 1 and 3 of the task, the first block of learning a given schema) and the second blocks (i.e., block 2 and 4 of the task, the second block of learning a given schema). A calculation of the difference between those two averages for first blocks (anticipatory-period average minus post-event boundary average) and the same for the second blocks was entered into a paired samples t-test for these two pre to post difference values.

The Shapiro-Wilk test was used to determine whether the assumption of normality was violated. If so, a Wilcoxon-signed rank test was used instead of a paired t-test. Effect sizes were calculated using Cohen’s d for t-tests and rank biserial correlation for wilxocon-signed rank tests. To visualize the univariate activity timelines around event boundaries, an extended period of TR = -10 up until TR = +10 was used for Figures 2 and 3 and for the Appendix Figures A3, A4, A5, A6 and A7. All statistical analysis were done using scipy (Virtanen et al., 2020) (as well as numpy, Harris et al., 2020, and pandas, McKinney, 2010, for data handling) and all visualizations were created using a combination between seaborn (Waskom, 2021) and matplotlib (Hunter, 2007).

**Fig. 2.**
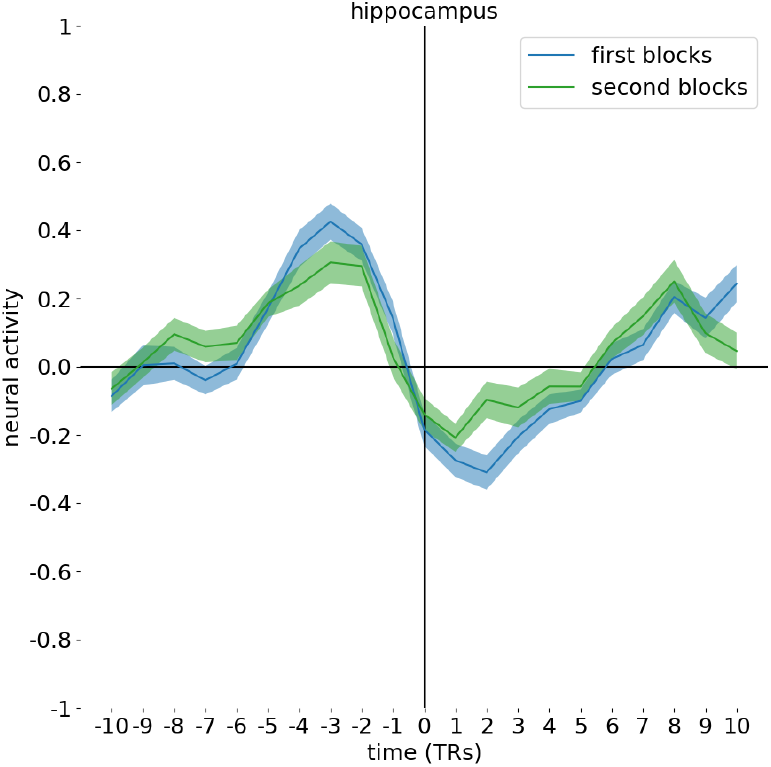
Univariate neural activity around event boundaries in the entire hippocampus (mean ± S.E.M.), with blue being the timeline of neural activity averaging both first blocks of schema learning –block 1 and 3– and green being the timeline averaging both second blocks –blocks 2 and 4– of schema learning. Time-point 0 is the event boundary.

**Fig. 3.**
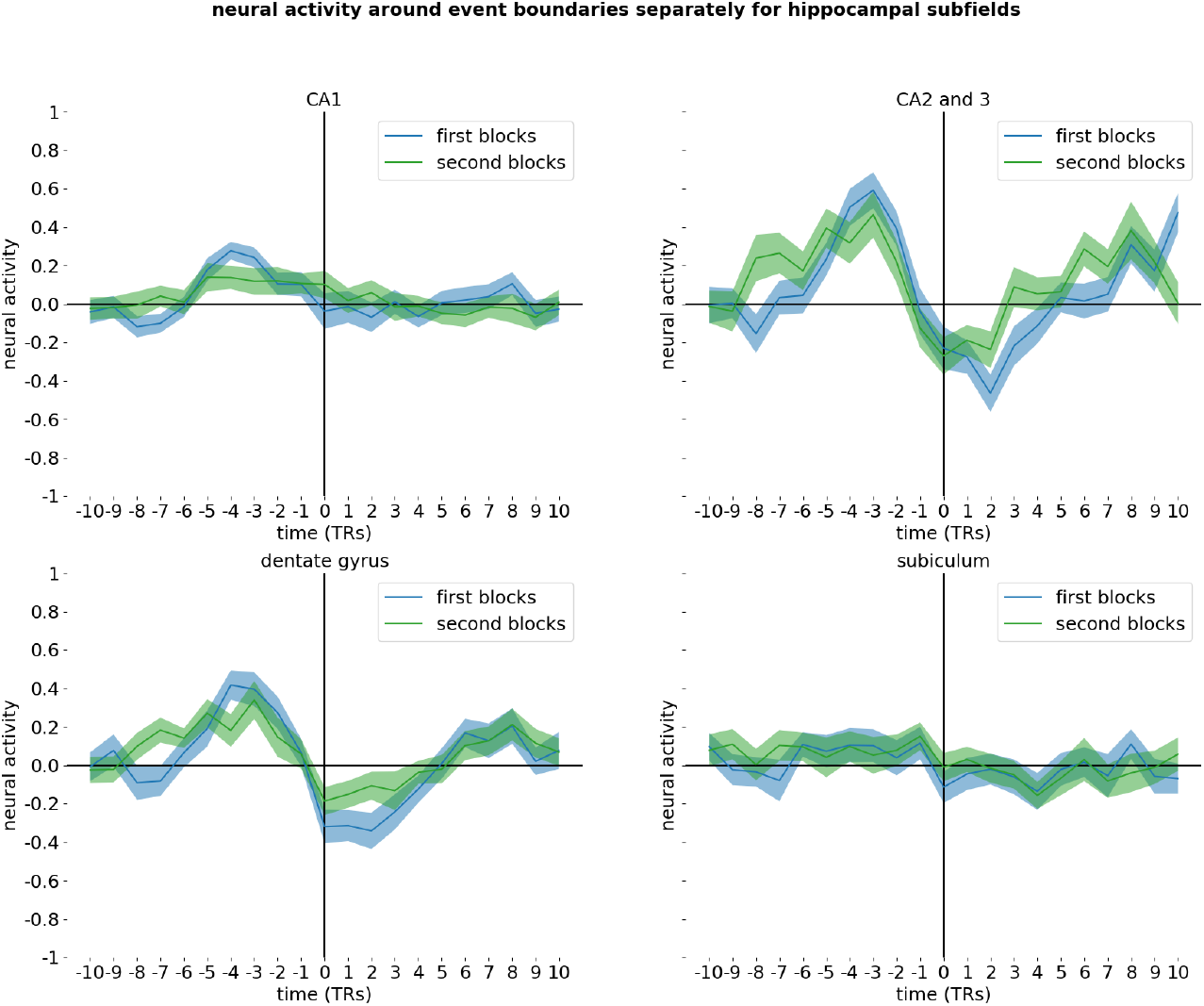
Univariate neural activity around event boundaries in the hippocampal subfields CA1, CA2/3, dentate gyrus and subiculum (mean ± S.E.M.), with blue being the timeline of neural activity averaging both first blocks of schema learning –block 1 and 3– and green being the timeline averaging both second blocks –blocks 2 and 4– of schema learning. Time-point 0 is the event boundary.

## 3 Results

### 3.1 Pre-boundary response: Hippocampal activity in anticipation to event boundaries

An average timeline of univariate hippocampal activity around event boundaries was calculated to investigate whether the hippocampus showed increased neural activation in anticipation to event boundaries. As shown in figure 2 and Appendix Figure A1, the results revealed that the hippocampal activity indeed peaks prior to the event boundaries, i.e., when the participant is in anticipation of an upcoming event boundary, as evidenced by a paired samples T-test showing stronger neural activity for the average of the 5 TRs before an event boundary compared to the average of the 5 TRs after event boundary (W = 39, *p <* 0.001, r = -0.79).

### 3.2 Pre-boundary hippocampal response is modulated by schema learning

Furthermore, the hippocampus showed evidence for diminished peaks of activity in anticipation of event boundaries because of schema learning, as evidenced by a significantly higher magnitude of peak in the first learning blocks –blocks 1 and 3– compared to second learning blocks –blocks 2 and 4– (T(39) = 4.14, *p <* 0.001, d = 0.65). Appendix Figure A3 shows data from each block of the task separately (i.e., block 1, 2, 3 and 4).

### 3.3 Hippocampal subfields

Next, potential *differences across hippocampal subfields* in anticipation of event boundaries were examined and related to schema learning. Having established that the hippocampus in general indeed showed the expected peak response in anticipation to event boundaries as well as responded stronger during first then second blocks, the same average timelines of neural activity in anticipation to event boundaries was calculated for the various hippocampal subfields. This allowed us to evaluate which hippocampal subfields (CA1, CA2/3, dentate gyrus, subiculum) were showing peaks of activity in anticipation to event boundaries, and subsequently which hippocampal subfields, if any, responded stronger in the beginning (i.e., first blocks) compared to the end (i.e., second blocks) of learning a new schema.

#### 3.3.1 CA1

The CA1 (Figure 3, top left; and Appendix Figure A2, top left) showed peaks of activity in anticipation to event boundaries (W = 122, *p <* 0.001, r = -0.61). Additionally, CA1 showed stronger peaks of activity in first blocks compared to second blocks (T(39) = 2.13, p = 0.039, d = 0.34). This would indicate that the amplitude of the CA1-peak in anticipation to event boundaries is sensitive to schema learning. Appendix Figure A4 shows data from each block of the task separately (i.e., run 1, 2, 3 and 4).

#### 3.3.2 CA2/3

The CA2/3 (Figure 3, top right; and Appendix Figure A2, top right) showed peaks of activity in anticipation to event boundaries (W = 68, *p <* 0.001, r = -0.73). Additionally, CA2/3 showed stronger peaks of activity in first blocks compared to second blocks (T(39) = 2.45, p = 0.019, d = 0.39). This would indicate that the amplitude of the CA2/3-peak in anticipation to event boundaries is sensitive to schema learning. Appendix Figure A5 shows data from each block of the task separately (i.e., run 1, 2, 3 and 4).

#### 3.3.3 dentate gyrus

The dentate gyrus (Figure 3, bottom left; and Appendix Figure A2, bottom left) showed peaks of activity in anticipation to event boundaries (T(39) = 6.30, *p <* 0.001, d = 0.996). Additionally, dentate gyrus showed stronger peaks of activity in first blocks compared to second blocks (T(39) = 2.47, p = 0.018, d = 0.39). This would indicate that the amplitude of the dentate gyrus-peak in anticipation to event boundaries is sensitive to schema learning. Appendix Figure A6 shows data from each block of the task separately (i.e., run 1, 2, 3 and 4).

#### 3.3.4 subiculum

The subiculum (Figure 3, bottom right; and Appendix Figure A2, bottom right) showed peaks of activity in anticipation to event boundaries (W = 203, p = 0.005, r = -0.44). However, the subiculum did not show a difference in amplitude of the neural activity in first blocks compared to second blocks (W = 402, p = 0.92, r = -0.017). This would indicate that the amplitude of the subiculum-peak in anticipation to event boundaries is *not* sensitive to schema learning. Appendix Figure A7 shows data from each block of the task separately (i.e., run 1, 2, 3 and 4).

Altogether, these results indicate that all hippocampal subfields have increased activity before an event boundary. Looking at the effect sizes, these pre-boundary responses are stronger for CA1, CA2/3 and dentate gyrus (with all effect sizes in the large effect range, i.e., 0.6-0.9), while the subiculum showed a moderate effect (i.e., an effect size around 0.4). Additionally, the results indicate that the CA1, CA2/3 and dentate gyrus showed stronger pre-boundary responses for blocks in the beginning of learning a new event schema compared to blocks at the end of learning a new event schema, indicating that these pre-boundary responses in those subfields are sensitive to schema learning (all with effect sizes in the moderate range, around 0.3/0.4). The subiculum did not show such a learning effect.

## 4 Discussion

The aim of this study was to investigate how event schema learning influences neural activity in anticipation to event boundaries across distinct hippocampal subfields during a naturalistic schema learning task. The results reveal that all subfields (i.e., CA1, CA2/3, dentate gyrus, and subiculum) are sensitive to event boundaries, as evidenced by increased activity in anticipation to an event boundary. Interestingly, it was only the CA1, CA2/3 and the dentate gyrus –and not the subiculum– that is also sensitive to schema learning, as evidenced by higher amplitude of peak activity in anticipation to event boundaries in the beginning of schema learning (i.e., when participants are still uncertain about upcoming events) compared to the end of schema learning (i.e., when participants can predict upcoming events better due to schema learning).

Learning a novel event schema happens by being exposed to many similar experiences drawn from that new event schema. Processing of all these experiences is accompanied by neural responses around boundaries between the events within these experiences. In particular the hippocampus has been suggested to be involved in “marking” these event boundaries by peaks in activity exactly at such event boundaries (Ben-Yakov & Henson, 2018; Bilkey & Jensen, 2021; Brunec et al., 2018; De Soares et al., 2024; Reagh et al., 2020). These hippocampal event boundary responses tend to reduce based on the strength of the prediction errors at those event boundaries (Ben-Yakov & Henson, 2018). In the current study, it was confirmed that the hippocampus is indeed sensitive to event boundaries, by showing a hippocampal peak of activity shortly before an event boundary. In contrast to the literature cited above, the current study showed the peak of activity to be present several seconds before the actual event boundary, rather than at or shortly after the event boundary. This indicates that the peak of activity happens when the participant is in anticipation of that event boundary, i.e., presumably when there is no prediction error yet. This could relate to how event boundaries were defined in this study (i.e., the objective switch between the event videos of a given wedding). In contrast to this, it is possible that participants subjectively experienced the event boundaries slightly earlier than the objective switch to the next event of the wedding. A plausible additional interpretation of this is that the peak of activity signals a moment of uncertainty about what will happen next, when the participant is expecting a new event to happen but is at that moment still uncertain about which one it will be. The fact that these peaks of activity reduced over the course of schema learning is in line with this interpretation. The last blocks of learning a new event schema –when participants are less and less uncertain about what event will happen next– showed reduced peaks in anticipation to event boundaries compared to the start of learning (when upcoming events are still uncertain for a participant). This is in line with the results from (Michelmann et al., 2021) that showed that hippocampal responses due to event boundaries shift in time, i.e., happen earlier once a participant can predict upcoming events better.

Here, possible differences between the various hippocampal subfields were investigated to get a more precise understanding of hippocampal activity in anticipation to event boundaries. The results showed increased neural activity in anticipation to an event boundary in CA1, CA2/3, dentate gyrus, and subiculum. However, the peak of activity pre-boundary in subiculum was much less pronounced then in the other subfields. Given that this task is focused on new learning and thus memory encoding, it was expected that the subiculum would not be involved as much as the other subfields (Seok & Cheong, 2020). CA1, CA2/3 and dentate gyrus all showed a decreased peak in anticipation to event boundaries at the end of schema learning (i.e., once the event sequences are highly predictable to participants), while the subiculum did not show this effect. Although CA1, CA2/3 and dentate gyrus all showed this effect, it was in particular the CA1 that seemed to only showed peaks at the beginning of schema learning and hardly any peak at all anymore at the end of schema learning. This is indicative of CA1 responses to event boundaries being influenced strongly by schema learning. CA1 is known to be involved in memory integration (Dimsdale-Zucker et al., 2018; Farovik et al., 2010; Molitor et al., 2021). Thus, schema learning might have led to participants integrating these various reoccurring types of events in memory. As a result of that, CA1 might have integrated these types of events in memory, which might indirectly have led to less surprise at event boundaries and therefore less strong of a pre-boundary response in the CA1. Animal work also suggests CA1 to be particularly responsive to novelty (Priestley et al., 2022) which would be in line with the complete absence of a peak in CA1 once the event sequences are not novel any longer.

Dentate gyrus showed sensitivity to event boundaries during the entire task, i.e., there is a clear peak in activity in the beginning as well as towards the end of the task. Somewhat surprisingly, it did also show sensitivity to schema learning, given that the activity peak was reduced at the end of schema learning compared to the beginning. Also when looking across each task-run individually, the dentate gyrus still showed a clear peak of activity in anticipation of event boundaries in the very last task block. This is distinct from the CA1 response, for which the peak of activity in anticipation to event boundaries is almost completely gone by the last task block. This is in line with the view that dentate gyrus is involved in separating similar information into distinct representations (Baker et al., 2016; Guo et al., 2025). The somewhat surprising effect that dentate gyrus peak responses in anticipation to event boundaries attenuate over the course of schema learning, might be in line though with some dentate gyrus theories that suggest that the dentate gyrus, although primarily focused on pattern separation, is also in some situations implicated in binding by associating incoming sensory information (Borzello et al., 2023).

The CA2/3 subfield showed sensitivity to anticipating the event boundaries as well as sensitivity to learning of the event schemas. CA3 has been suggested to be involved in separating similar information (Leal et al., 2014; Molitor et al., 2021) as well as integrating information (Bein & Davachi, 2024; Dimsdale-Zucker et al., 2018). A possible reason for the discrepancy between those results can be found in animal place cell research during condition experiments. S. Leutgeb et al., 2004 suggests that CA3 cells shows activity patterns in line with pattern separation at first, but switches to pattern completion eventually and only when input similarity increased. This suggests that CA3 can be used flexibly to both differentiate between similar information but also to integrate related information. Also Guo et al., 2025 suggests that CA3 would create distinct representations of highly similar information. Results of the current study would be in line with the CA3 integrating related information over the course of schema learning in this task, which would lead to diminished peaks of activity in anticipation to event boundaries at the end of schema learning (i.e., when the event sequences are known to participants).

It would be interesting to relate these findings to recall performance or retrieval processes in general. Especially given the fact that the various hippocampal subfields were shown to be either primarily involved with memory encoding or memory retrieval, results might be different in a more memory retrieval focused task. Additionally, it is of importance to translate these findings to children. While the task used here did mimic new schema learning in adults, most new schema learning in real life happens in children. Children are still, much more than adults, exploring the world and learning what to expect in which environment. Importantly, their hippocampus and prefrontal cortex, both regions important in schema learning, are still in development. Therefore, neural activity changes for emerging schemas might be different in children.

Altogether, results discovered by the current study are building on established findings about hippocampal responses to event boundaries and their relation to prediction errors by providing a more precise understanding about how the various hippocampal subfields will show anticipation to event boundaries, and furthermore how they adapt over the course of initial learning of novel event schemas.

## Data and code availability statement

The raw data and code will be made available upon publication of the article.

## Acknowledgements

I want to thank Prof. Kenneth Norman for helpful advise on data analysis of this project and for feedback on an earlier version of this manuscript. This work was supported by a Multi-University Research Initiative Grant (ONR/DoDN00014-17-1-2961) and an NWO Rubicon grant (446-17-009).

## Appendix A

**Fig. A1.**
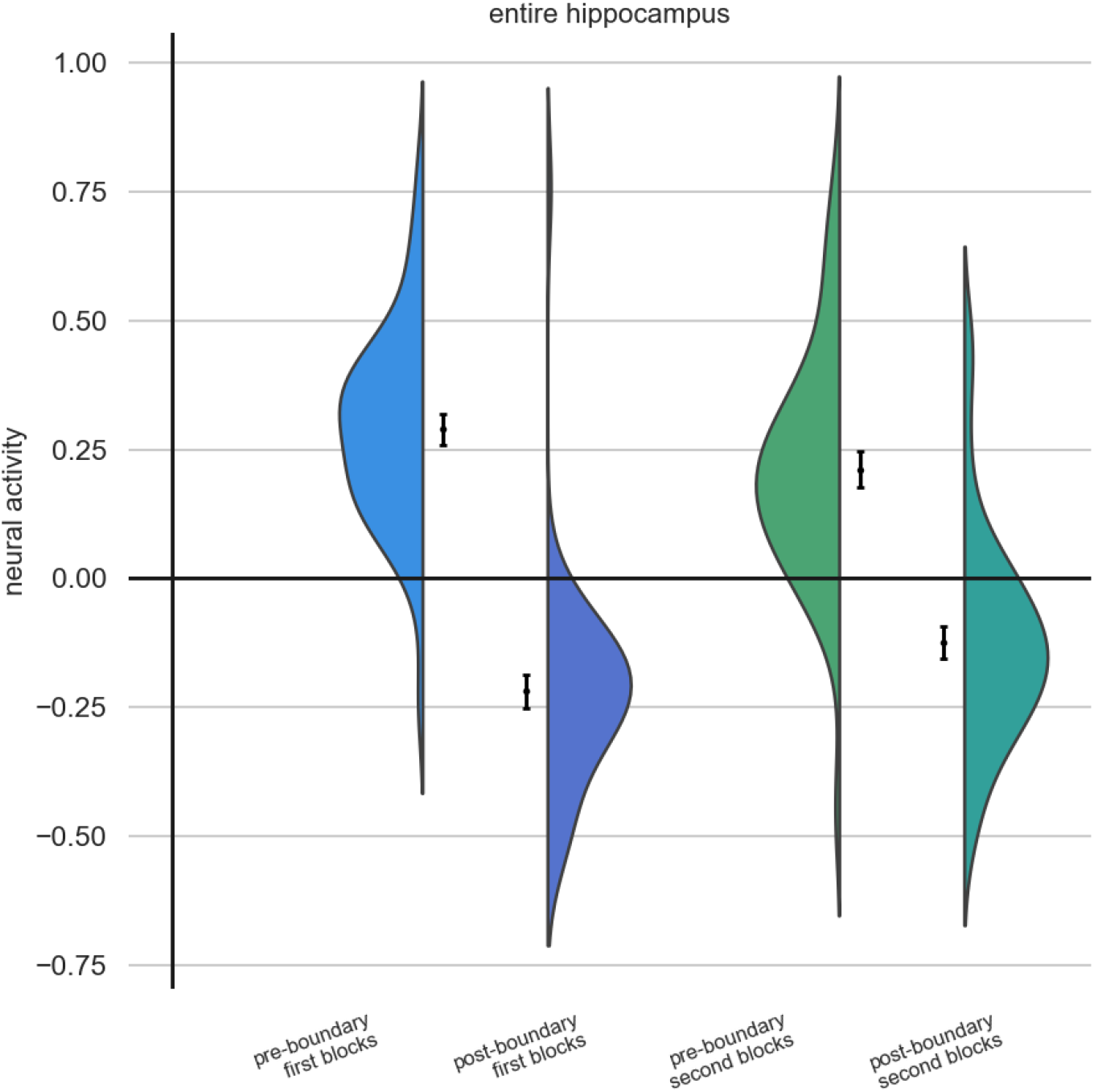
Average univariate neural activity pre-boundary (i.e., TR = -5 to TR = -1) and average univariate neural activity post-boundary (i.e., TR = 0 to TR = 4) in the entire hippocampus. Data is represented both as a violin plot to illustrate the distribution of the data and as mean ± S.E.M. The two blue bars represent the data pre- and post-event boundary for first blocks, and the green bars represent the data pre- and post-event boundary for second blocks.

**Fig. A2.**
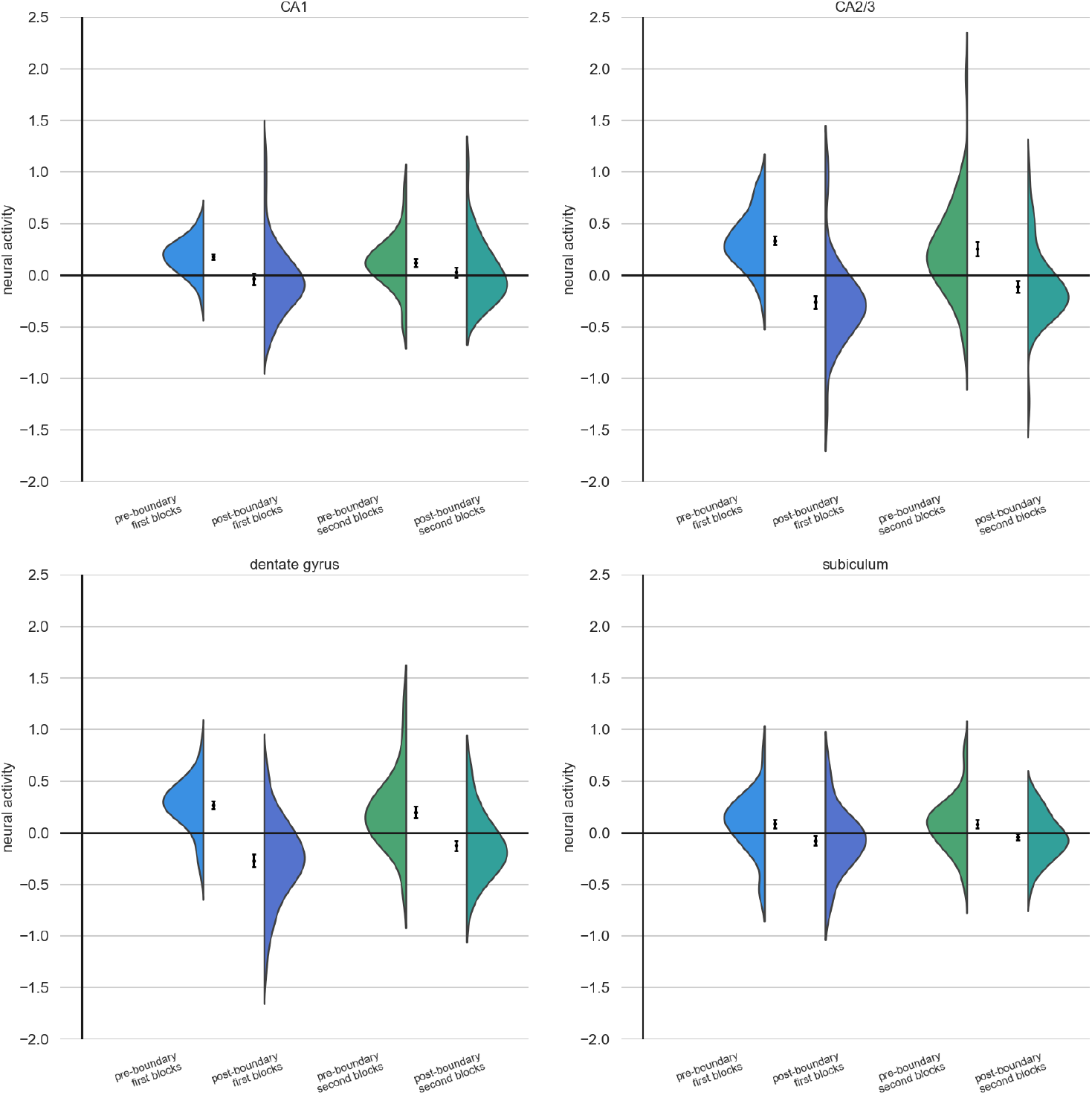
Average univariate neural activity pre-boundary (i.e., TR = -5 to TR = -1) and average univariate neural activity post-boundary (i.e., TR = 0 to TR = 4), separately for the CA1, CA2/3, dentate gyrus and subiculum. Data is represented both as a violin plot to illustrate the distribution of the data and as mean ± S.E.M. For each subplot, the two blue bars represent the data pre- and post-event boundary for first blocks, and the green bars represent the data pre- and post-event boundary for second blocks.

**Fig. A3.**
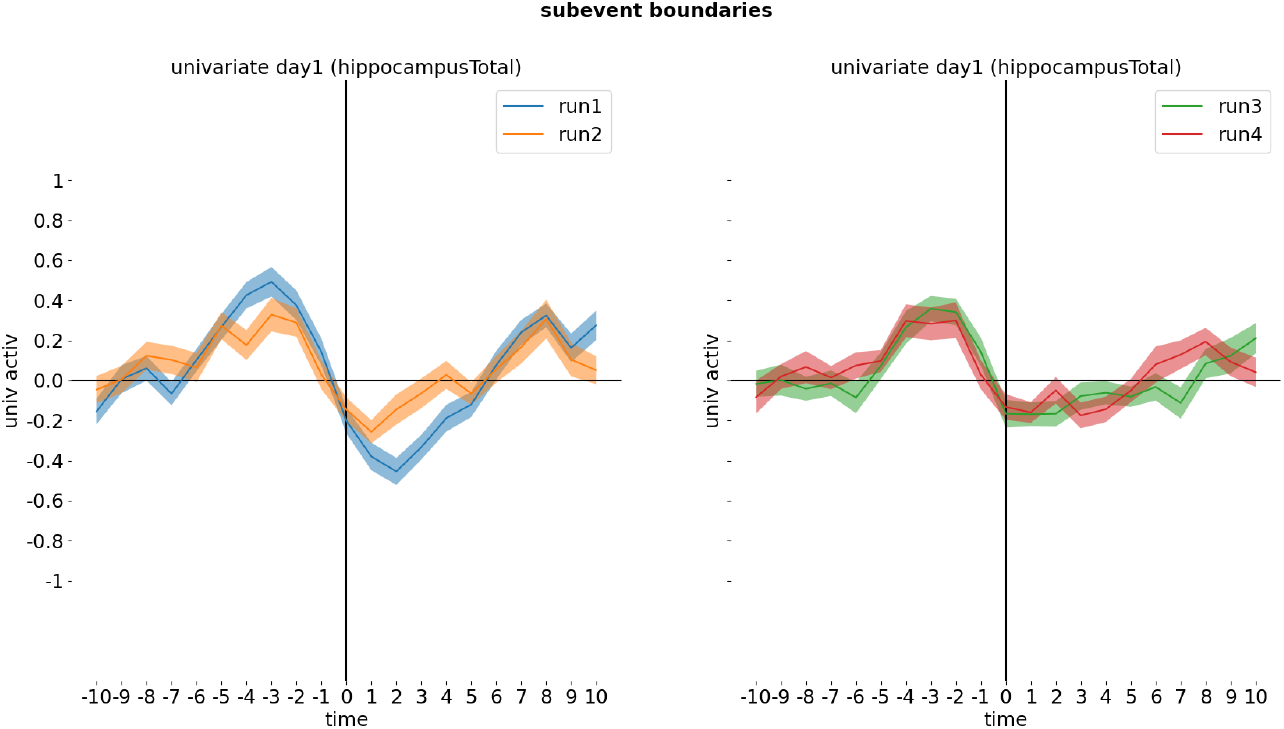
Univariate neural activity around event boundaries in the entire hippocampus (mean ± S.E.M.), separately for block 1 (blue), 2 (orange), 3 (green) and 4 (red).

**Fig. A4.**
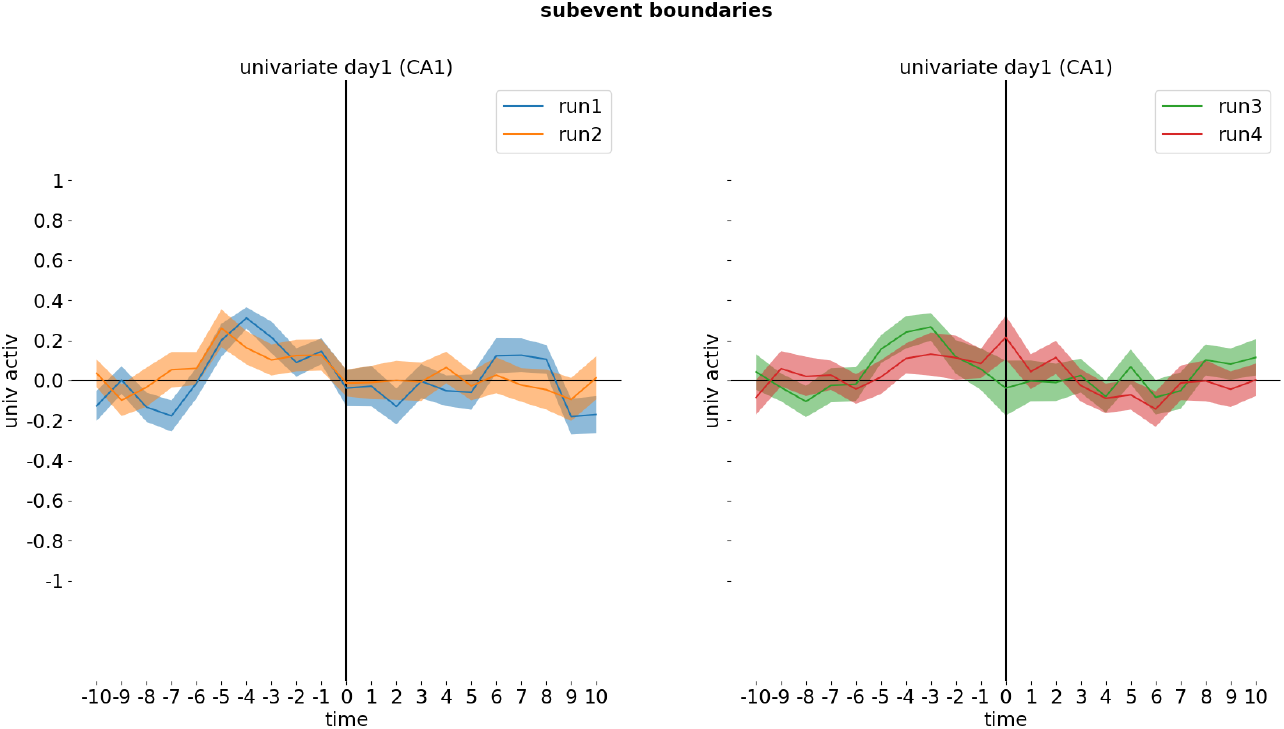
Univariate neural activity around event boundaries in the CA1 (mean ± S.E.M.), separately for block 1 (blue), 2 (orange), 3 (green) and 4 (red).

**Fig. A5.**
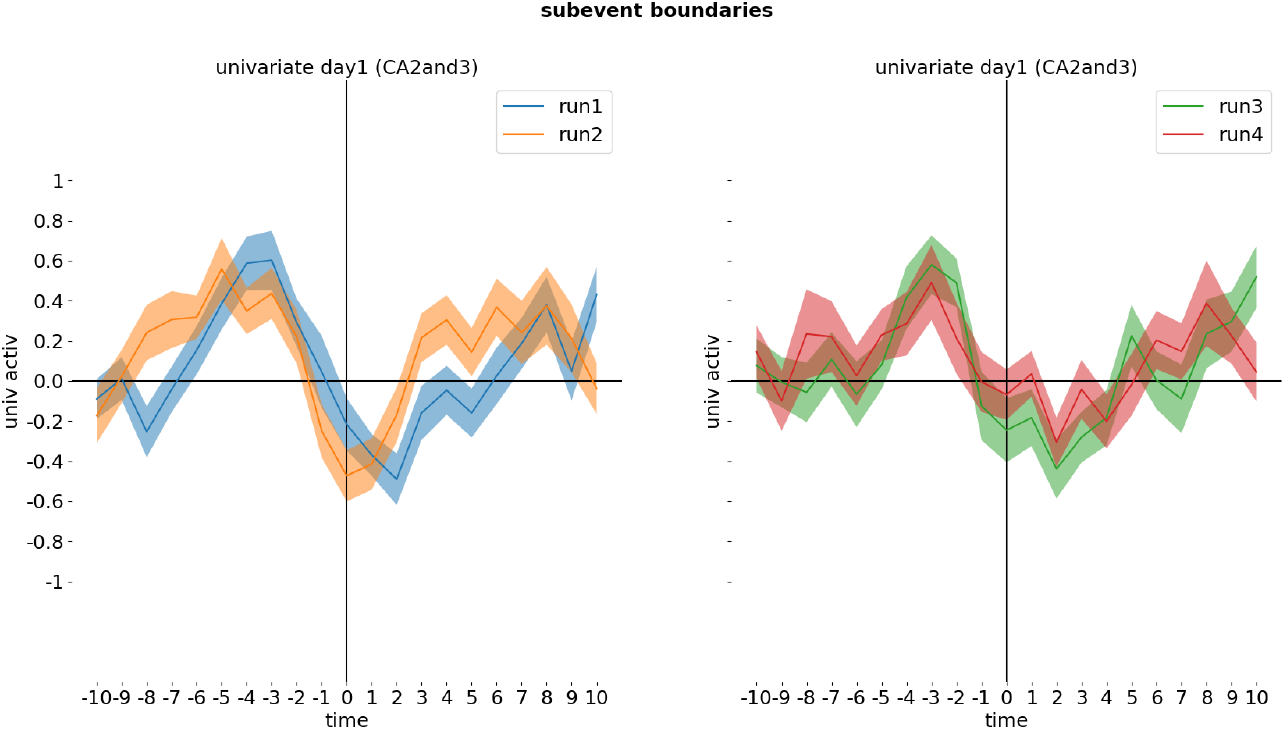
Univariate neural activity around event boundaries in the CA2/3 (mean ± S.E.M.), separately for block 1 (blue), 2 (orange), 3 (green) and 4 (red).

**Fig. A6.**
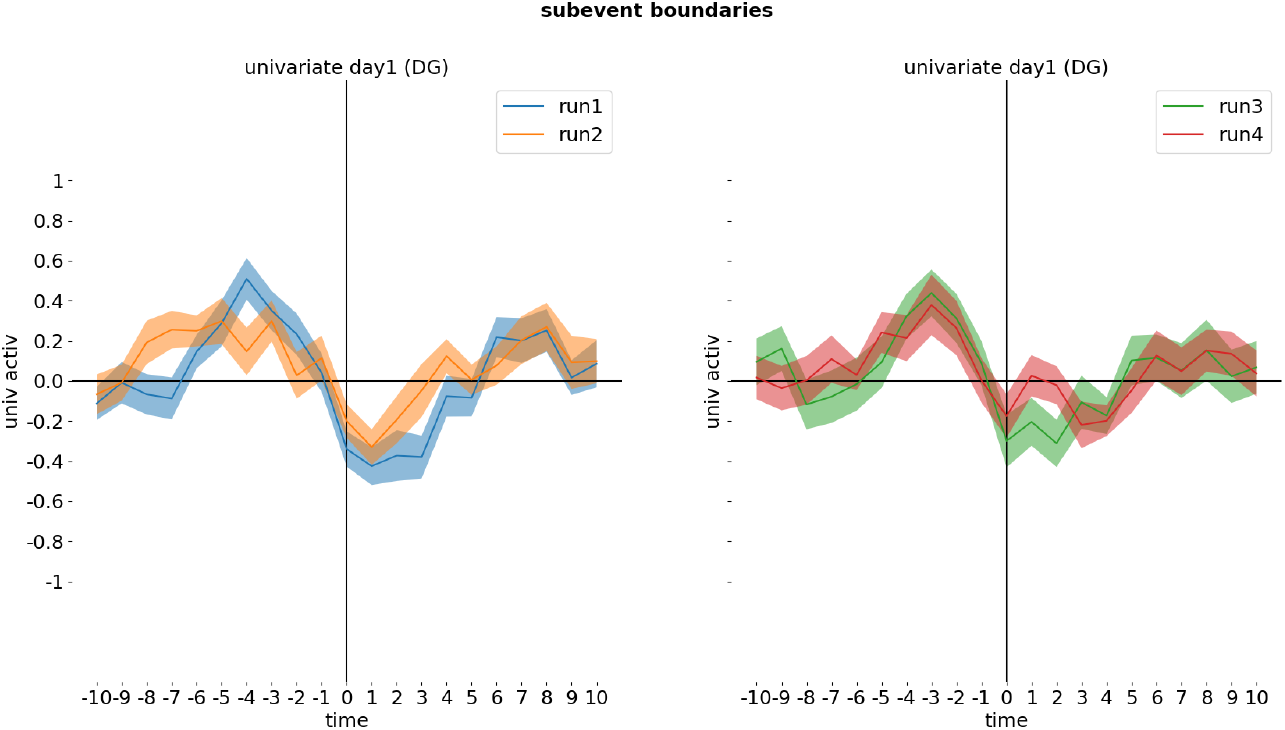
Univariate neural activity around event boundaries in the dentate gyrus (mean ± S.E.M.), separately for block 1 (blue), 2 (orange), 3 (green) and 4 (red).

**Fig. A7.**
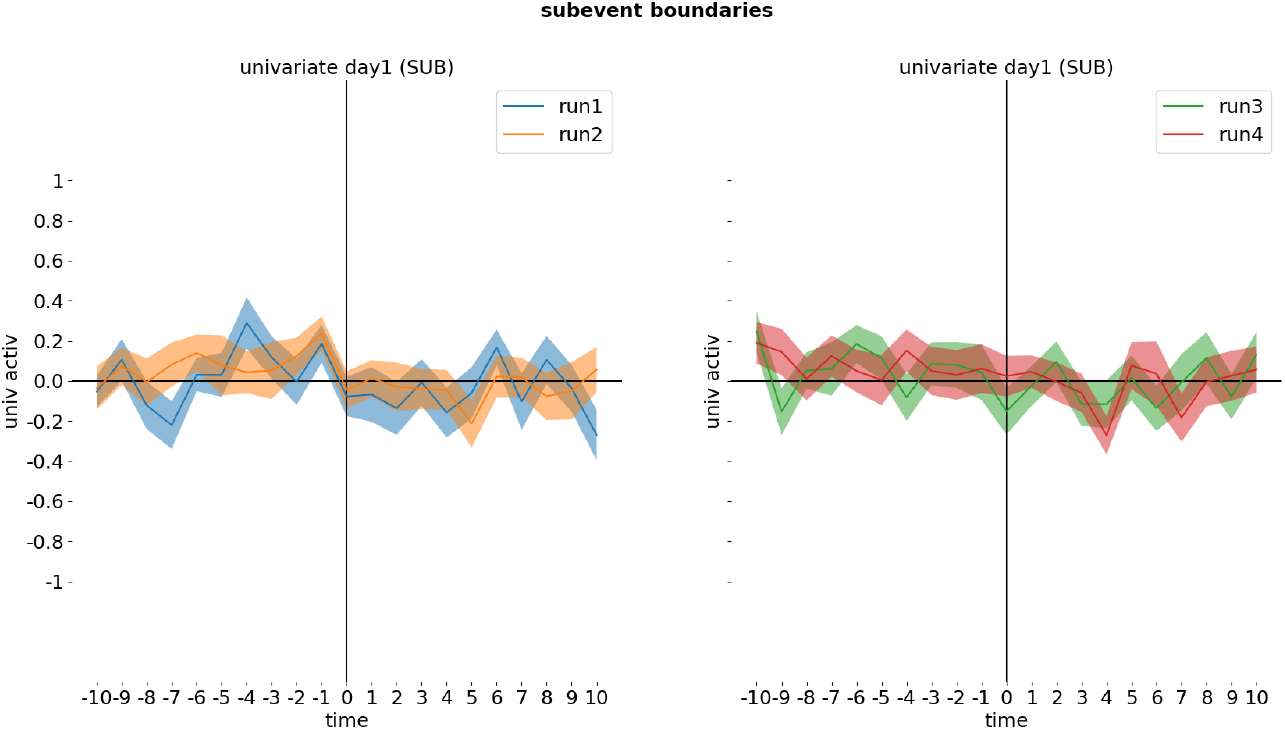
Univariate neural activity around event boundaries in the subiculum (mean ± S.E.M.), separately for block 1 (blue), 2 (orange), 3 (green) and 4 (red).

